# Structure of human hephaestin reveals a homodimer anchored to the cell membrane

**DOI:** 10.64898/2026.11.09.765449

**Authors:** Ce Liang, Ming Zhou, Yaping Pan

## Abstract

Hephaestin (Heph) is a membrane-bound ferroxidase that oxidizes ferrous ion (Fe^2+^) exported by ferroportin (Fpn) to ferric ion (Fe^3+^), enabling iron loading onto transferrin for systemic delivery. Loss-of-function mutations in Heph cause iron accumulation in intestinal enterocytes and systemic iron deficiency in animal models, underscoring its role in iron export. Heph has a large extracellular domain anchored to the cell membrane by a single transmembrane helix. However, full-length Heph has not previously been purified, and our understanding of its structure and function remains incomplete. Here, we report the expression and purification of functional full- length human Heph and determination of its structure by cryo-electron microscopy at an overall resolution of approximately 2.3 Å. The structure reveals a stable homodimer with well-resolved extracellular domains. Although the transmembrane helices are less well resolved, their positions can be defined. The substrate-binding site is connected to an electronegative funnel that may facilitate Fe^2+^ capture. These findings establish the structural organization of full-length human Heph and provide structural constraints for understanding its interaction with Fpn.

## Introduction

Hephaestin (Heph), encoded by the *HEPH* gene, is a membrane-bound ferroxidase expressed predominantly in the small intestine^1^. Heph is known to partner with ferroportin (Fpn)^2–5^, which is an iron exporter, and oxidizes ferrous ions (Fe^2+^) to ferric ions (Fe^3+^). Conversion of Fe^2+^ to Fe^3+^ is essential because it helps maintain the driving force for Fe^2+^ export, prevents harmful reactive oxygen species produced by Fe^2+^, and enables loading of transferrin, which recognizes only the ferric ions^6,7^. Loss of Heph activity leads to impaired iron export and accumulation of iron in enterocytes and anemia, as seen in sex-linked anemia (*sla*) mice that carry a large in- frame deletion in Heph^1^. Similar phenotypes were also reported in Heph-knockout (*HEPH*^-/-^) mice^8,9^, which re-enforces the role of Heph in iron absorption and homeostasis.

Heph is composed of an N-terminal extracellular domain, ∼1100 amino acids, anchored to the plasma membrane by a single transmembrane helix, 34 amino acids. The protein sequence of the extracellular domain is ∼50% identical to that of ceruloplasmin, a secreted soluble ferroxidase^10,11^. While structure and function of ceruloplasmin have been characterized extensively^12–14^, studies of Heph remain limited. The extracellular domain of Heph has been expressed in previous studies and shown catalytically competent^15–18^. Although the overall structural fold and potential metal ion binding sites in Heph can be inferred from ceruloplasmin, it remains unknown how the catalytic and substrate-binding regions of Heph are oriented relative to the membrane, which is important for understanding its potential interaction with Fpn and the capture of Fe^2+^. Structural analysis of full-length Heph is therefore needed to define its membrane-associated organization.

In the current study, we expressed and purified full-length human Heph from HEK293 cells, demonstrated ferroxidase activity of the purified Heph, and determined its structure by cryo- electron microscopy (cryo-EM).

## Results

### Recombinant Heph retains ferroxidase activity

Full-length human Heph (UniProt ID: Q9BQS7) was expressed in HEK293 cells and purified (**Figure 1A, B**). A characteristic absorbance peak at ∼610 nm on the UV-Vis spectrum is consistent with the presence of type 1 copper centers^19,20^ (**Figure 1C**). Purified Heph protein shows ferroxidase activity in a ferrozine-based assay^15^ (**Figure 1D**). Oxidation of Fe^2+^ displays saturable kinetics, with an apparent *Km* of 8.1 µM and a *Vmax* of 10.9 µM min⁻¹ (**Figure 1E**). Pre- treatment of Heph with 1 mM D-penicillamine, a copper chelator, suppressed ferroxidase activity. Together, these results demonstrate that the purified full-length Heph contains copper cofactors and retains ferroxidase activity.

**Figure 1.**
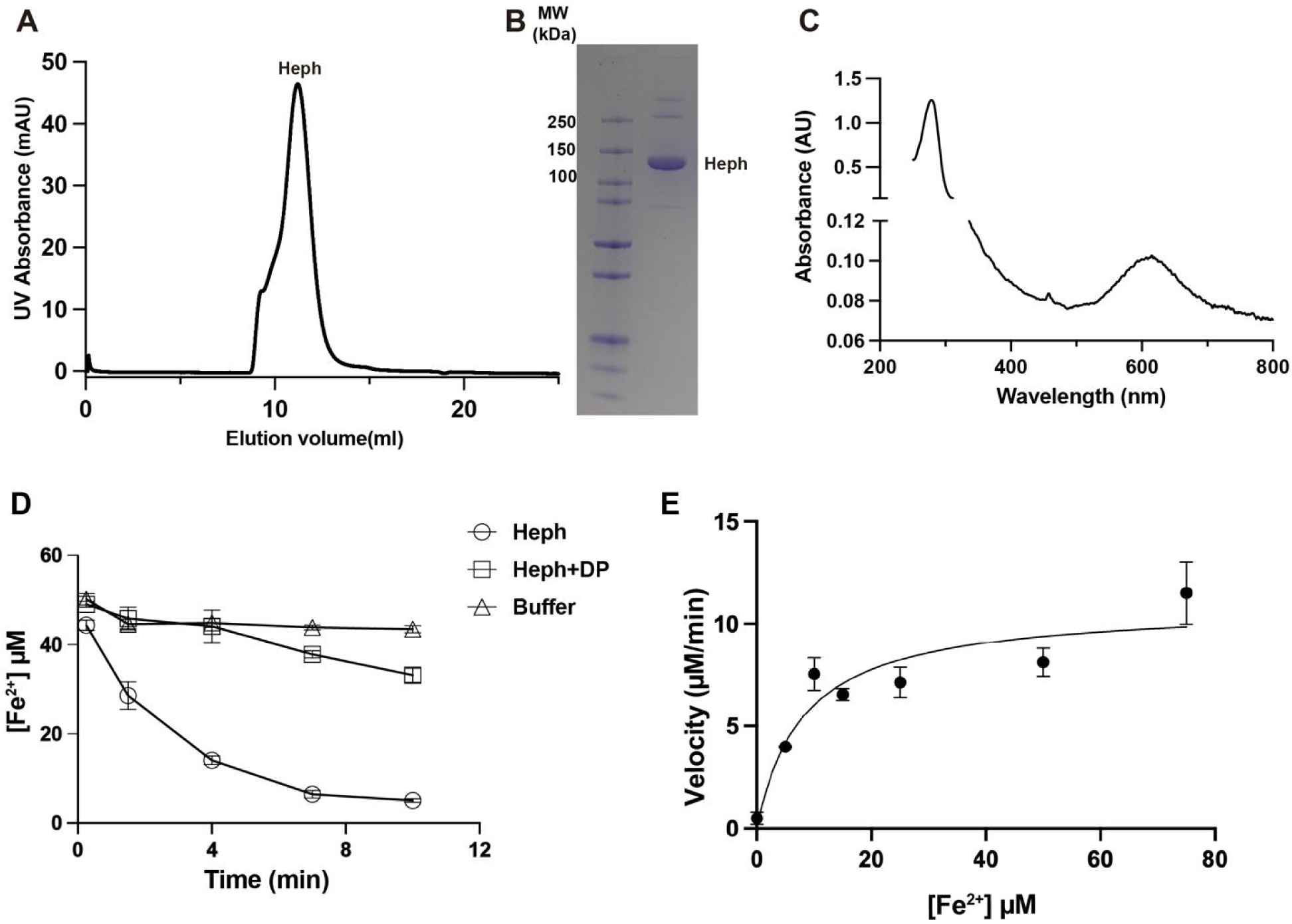
Purification and ferroxidase activity of full-length human Heph. (A) Size- exclusion chromatography profile of purified full-length human Heph on a Superose 6 Increase 10/300 GL column. (B) SDS–PAGE analysis of purified Heph. Molecular mass markers are indicated in kDa. (C) UV-Vis absorption spectrum of purified Heph, showing an absorbance peak near 610 nm consistent with the presence of type 1 copper centers. (D) Time course of Fe²⁺ oxidation by untreated Heph (circles), Heph pretreated with the copper chelator D-penicillamine (DP; squares), or buffer alone (triangles). Reactions contained 50 µM Fe²⁺ and 3.6 µM Heph. Residual Fe²⁺ was quantified using a ferrozine assay. Free DP was removed by size-exclusion chromatography before activity measurements. (E) Initial reaction velocities measured with 0.2 µM Heph at varying Fe²⁺ concentrations. The data fit to the Michaelis–Menten equation, yielding an apparent K of 8.1 µM and a V ₐₓ of 10.9 µM min⁻¹. Assays in D and E were performed at 37 °C, and data are presented as mean ± SD (n = 3).

## Overall structure of Heph dimer

Structure of the full-length human Heph was determined by single-particle cryo-EM, with the extracellular domain resolved at ∼2.3 Å resolution and the transmembrane domain at ∼6-8 Å (**Figure 2A, Figure S1, Supplementary Table 1**).

**Figure 2.**
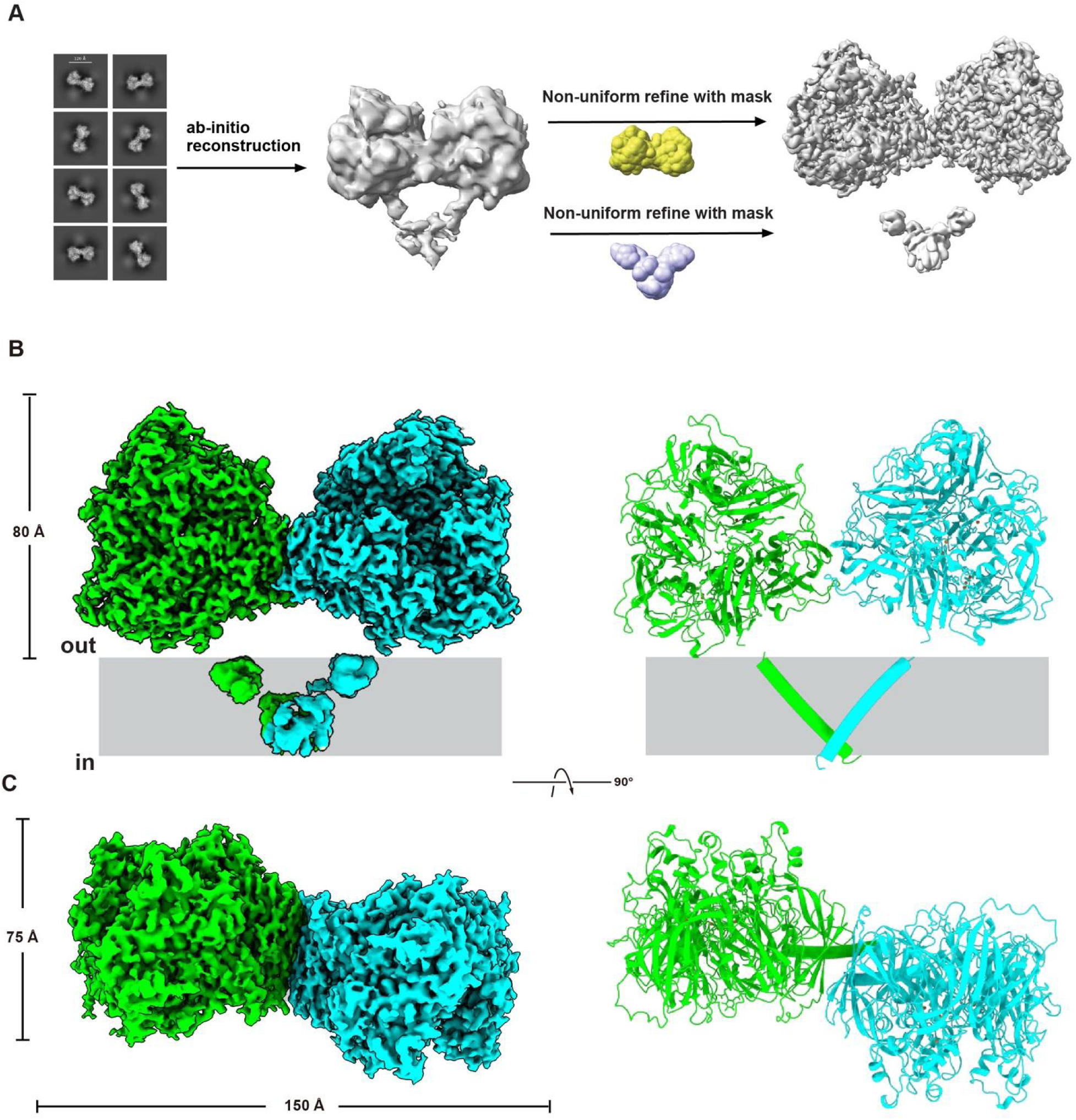
Cryo-EM structure of the full-length human Heph homodimer. (A) Overview of cryo-EM reconstruction and refinement. Representative two-dimensional class averages were used to generate an ab initio reconstruction. Separate nonuniform refinements using masks encompassing the extracellular region (yellow) or transmembrane region (purple) yielded maps at approximately 2.3 Å and 6–8 Å resolution, respectively. (B) Side views of the cryo-EM density maps (left) and structural model (right), with the two Heph protomers colored green and cyan. The extracellular domains are shown as ribbons, and the transmembrane helices are represented as cylinders docked into the lower-resolution density. The gray band indicates the membrane. (C) Orthogonal views of the density map (left) and structural model (right), rotated by 90° relative to B. The extracellular assembly measures approximately 150 Å × 75 Å in this view and extends approximately 80 Å above the membrane in B.

The entire extracellular domain of Heph, residues 23-1065, is resolved in the density map (**Figure 2B, C**). Two extracellular domains are welded together by a substantial dimer interface that has a buried surface area of 796 Å^2^. Some of the contacting residues at the dimer interface are highlighted in **Figure 3A-B**. The density map for the two TM domains is only sufficient for docking of two α-helices, and the two helices seem to cross over and make contacts towards the intracellular side of the membrane (**Figure 2B, C**). To test whether the homodimer is present on cell membrane, we used a split luciferase assay (NanoBiT)^21^, in which two complementary parts of a luciferase were individually spliced to the C-terminus of a Heph, Heph-SmBiT and Heph- LgBiT (**Figure 3C**). When the two constructs are co-expressed in HEK cells, robust luciferase activity was detected (**Figure 3D**), indicating assembly of the two Heph constructs.

**Figure 3.**
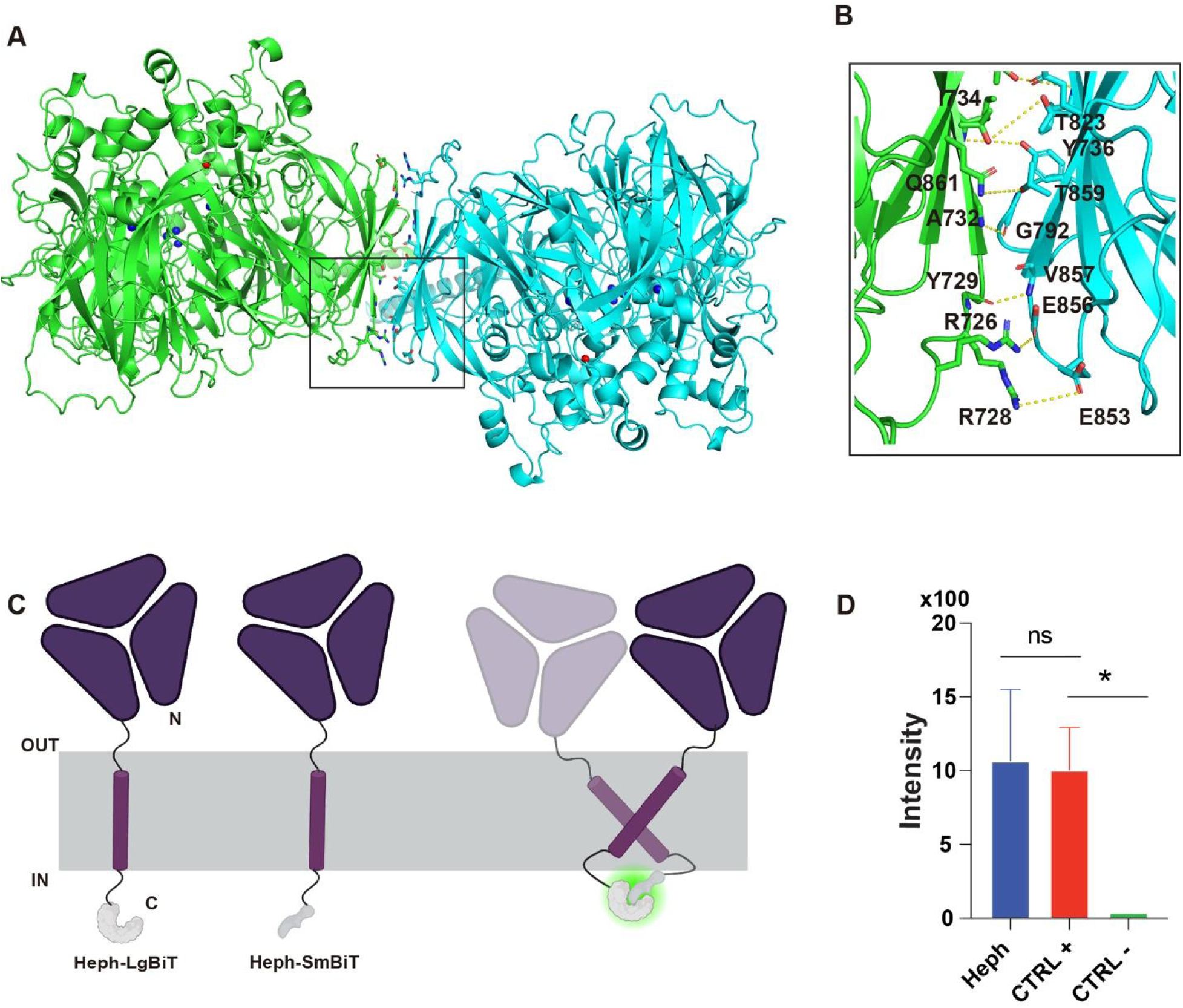
Dimer interface and cellular self-association of human Heph. (A) Ribbon representation of the Heph dimer, with the two protomers colored green and cyan. The boxed region highlights the extracellular dimer interface shown in B. (B) Close-up of the dimer interface. Selected interacting residues are shown as sticks and labeled; yellow dashed lines indicate putative polar interactions. (C) Schematic of the NanoBiT luciferase complementation assay. The large (LgBiT) and small (SmBiT) luciferase fragments were fused to the C-termini of Heph. Association of the fusion proteins brings the fragments into proximity, reconstituting luciferase activity. The gray band represents the membrane. (D) Luminescence signals from HEK cells coexpressing Heph-LgBiT and Heph-SmBiT (Heph), alongside positive (CTRL+) and negative (CTRL−) controls. Data are presented as mean ± SD (n = 3). P < 0.05 is indicated by an asterisk; ns, not significant.

## Structure of the extracellular domain

The structure of an individual Heph extracellular domain is similar to that of ceruloplasmin (PDB 1KCW)^12^, and the two can be superposed with a root mean squared distance (rmsd) of 2.0 Å across 1002 aligned Cα atoms (**Figure S2A**). Each extracellular domain consists of six consecutive cupredoxin-like domains (D1-D6) arranged sequentially around a central axis to form the characteristic multicopper oxidase fold (**Figure 4A**).

**Figure 4.**
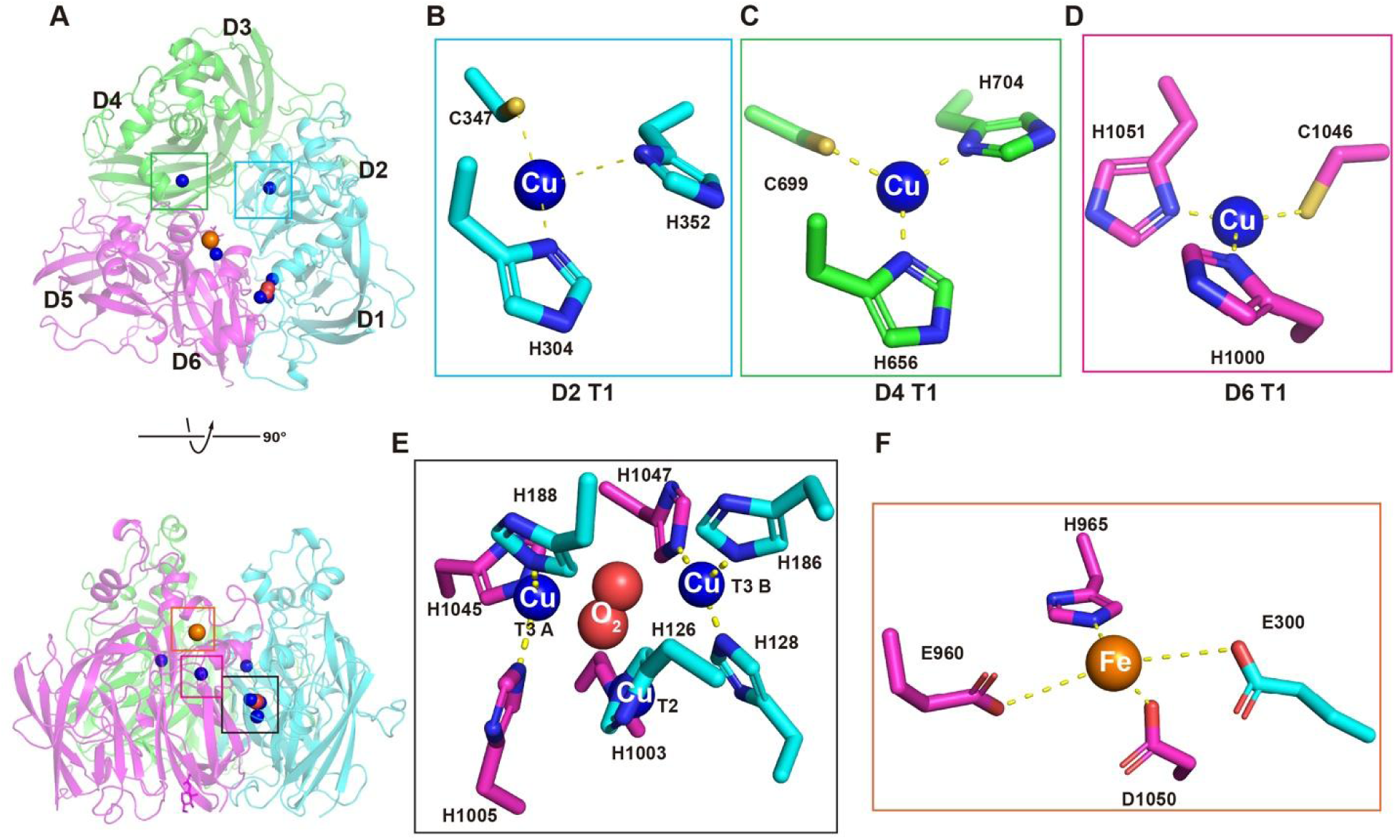
Domain organization and metal-binding sites of human Heph. (A) Ribbon representation of the extracellular region of one Heph protomer in two views related by a 90° rotation. The six cupredoxin-like domains are labeled D1–D6, with D1 and D2 colored cyan, D3 and D4 green, and D5 and D6 magenta. Copper and iron ions are shown as blue and orange spheres, respectively. Boxes indicate the regions enlarged in B–F. (B–D) Coordination of the type 1 (T1) copper centers in D2 (B), D4 (C), and D6 (D), showing selected coordinating residues. (E) Trinuclear copper center at the interface between D1 and D6, comprising one type 2 (T2) and two type 3 (T3A and T3B) copper ions. The O₂ molecule modeled within the center is shown as red spheres. (F) Iron-binding site adjacent to the D6 T1 copper center. The modeled iron ion is coordinated by E300, E960, H965, and D1050. In B–F, coordinating residues are shown as sticks and labeled, and yellow dashed lines indicate coordination of contacts.

The density map reveals six well-defined copper binding sites (**Figure 4A**), which are conserved between Heph and ceruloplasmin (**Figure S3**). Three mononuclear type 1 (T1) copper centers are found in domains 2, 4, and 6, each coordinated by H304/C347/H352, H656/C699/H704, and H1000/C1046/H1051, respectively (**Figure 4B-D**). The remaining three copper ion binding sites form the trinuclear center at the interface between D1 and D6, with a type 2 copper site (H126 and H1003), and two type 3 copper sites (H128/H186/H1047 and H188/H1005/H1045) (**Figure 4E**). An additional density is found in the center of the three Cu, in which an oxygen (O_2_) molecule was placed. The oxygen molecule was also observed in the structures of ceruloplasmin (PDBs 2J5W and 4ENZ)^14,22^, and is the final electron acceptor that sustains the ferroxidase activity. Both the location and the coordinating residues of the six copper binding sites are well- conserved between Heph and ceruloplasmin (**Figure S3**), with the exception of the D2 T1 copper site in which Heph has an additional M357 at the axial position of the copper ion (**Figure S2B**).

A single substrate (Fe^2+^) binding site is marked by a non-protein density adjacent to the D6 T1 copper site (**Figure 4F**), and the Fe^2+^ is coordinated by the side chains of E300/E960/H965/D1050. Previous mutational studies showed that E960 and H965 are critical for high-affinity Fe^2+^ binding and ferroxidase activity^18^. A similar substrate binding site is found in ceruloplasmin, which is coordinated by E272/E935/H940/D1025 (**Figure S3**).

## Discussion

In this study, we purified functional full-length human Heph and determined its structure by cryo-EM. The structure reveals an unexpected homodimeric assembly, which may reduce the mobility of the extracellular domain and enhance its encounter with Fpn.

Dimerization of Heph is mediated by both the extracellular and membrane-spanning regions. Results from the NanoBiT complementation assay provide independent evidence supporting Heph dimer in cell. This assay does not, however, establish its endogenous stoichiometry or distinguish cell-surface association from association in intracellular compartments. Furthermore, dimerization may enhance the stability or catalytic activity of Heph, which requires additional experiments by comparing the wild type Heph with properly folded mutants that disrupt dimerization.

The full-length Heph structure places the substrate binding site close to the membrane, providing a possible explanation for efficient capture of Fe^2+^ released by Fpn. Capture of Fe^2+^ released by Fpn is facilitated by two factors. First, with the extracellular domain being anchored to the cell membrane, the Fe^2+^ binding site on Heph is confined to within <20 Å of the membrane surface to reduce free diffusion of Fe^2+^ (**Figure 5A**). Second, electrostatic potentials on the surface of Heph show that the Fe^2+^ binding site is located at the center of a deeply electronegative well (**Figure 5A**), which helps funnel Fe^2+^ into the substrate binding site.

**Figure 5.**
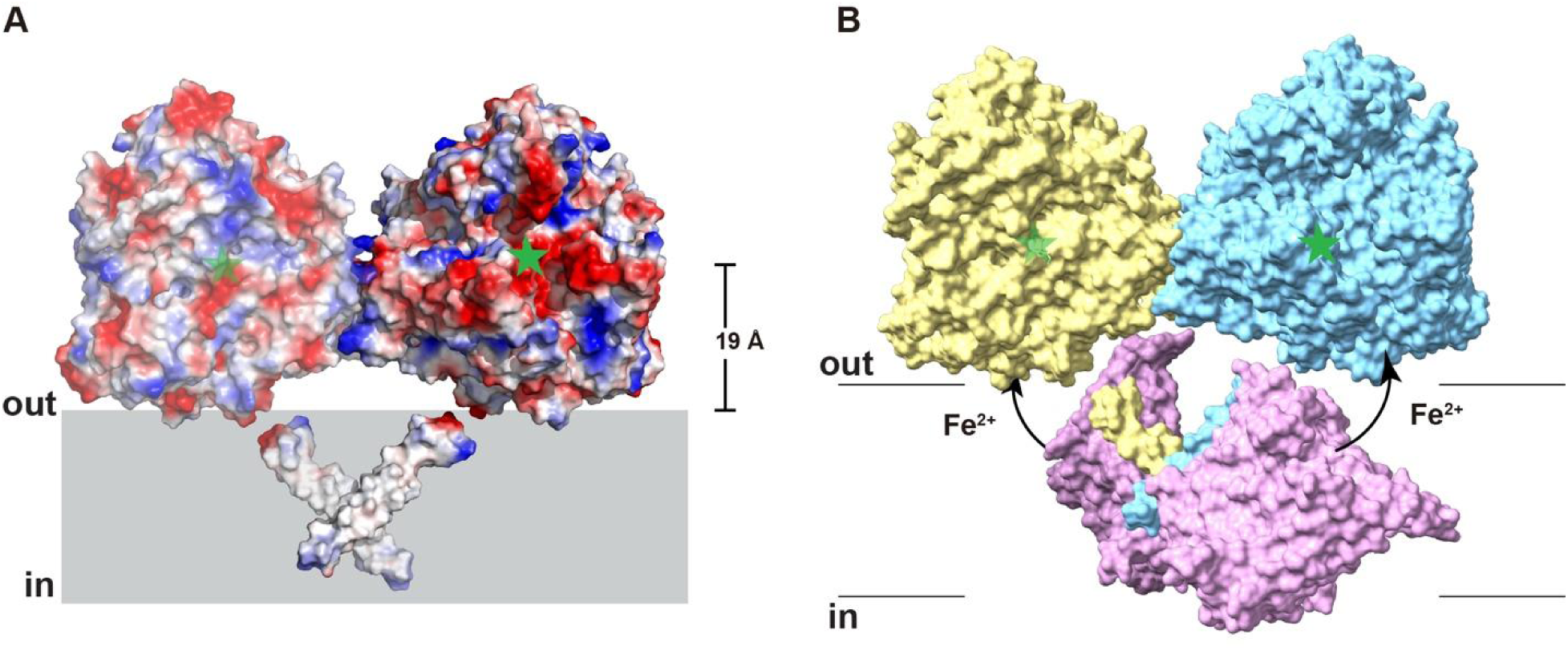
Membrane positioning of Heph and proposed coupling to Fpn. (A) Surface representation of the Heph dimer colored by electrostatic potential, with red, white, and blue indicating negative, neutral, and positive potentials, respectively. Green stars mark the Fe²⁺- binding sites within negatively charged surface pockets. The indicated distance places the substrate-binding site approximately 19 Å above the extracellular membrane surface in the illustrated membrane placement. (B) Proposed arrangement of a Heph dimer and two Fpn molecules. Heph protomers are colored yellow and cyan, and Fpn molecules are colored pink. Green stars mark the Fe²⁺-binding sites on oppositely facing surfaces of the Heph dimer. Arrows indicate proposed paths of Fe²⁺ transfer from each Fpn molecule toward the adjacent Heph substrate binding site. Horizontal lines indicate membrane boundaries.

Previous cellular studies reported association or close proximity between Heph and Fpn^2–5^, but a potential Heph-Fpn complex has not been defined. The structure of the Heph dimer imposes constraints to the assembly of Heph with Fpn. Since the two substrate binding sites face the opposite directions, it is likely that Fpn associates with the Heph dimer from the opposite side of the dimer as well, as illustrated in **Figure 5B**. Such an arrangement would imply that one Heph dimer could engage two Fpn transporters. A structure of the full-length Heph-Fpn complex will be needed to determine how transporter association is accommodated alongside Heph dimerization.

Overall, our study defines the structural basis of human Heph ferroxidase activity and reveals several features that distinguish it from the closely related soluble ferroxidase ceruloplasmin. The conserved multicopper centers provide the catalytic framework for Fe^2+^ oxidation, whereas the homodimeric organization, distinct D2 T1 coordination environment, and D6 substrate binding site highlight structural features specific to Heph. These findings enhance our understanding of Heph-mediated iron oxidation at the cell surface.

## Methods

### Cloning, expression, and purification of Heph

The human Heph gene (UniProt ID: Q9BQS7) was codon-optimized for human cell expression and subcloned into a BacMam expression vector^23^ with green fluorescent protein (GFP) tag attached to the C-terminus. A tobacco etch virus (TEV) protease recognition site was inserted between the C-terminus of the protein and the GFP tag. The resulting plasmid containing cDNA of Human Heph was used to generate baculoviruses following the manufacturer’s protocol (Invitrogen). HEK293S GnTI- suspension cells at 3×106 cells/mL in Freestyle 293 medium (GIBCO) supplemented with 2% FBS were infected with 10% P3 virus. 1 M CuSO4 was filtered and added to the cell culture to reach a final concentration of 50 µM. Infected cells were incubated at 37 °C overnight (∼16 h). The protein expression was then induced by adding 10 mM sodium butyrate and temperature was decreased to 30 °C for two days (∼48 h) before harvesting. Cell pellets were homogenized in buffer A containing 20 mM HEPES (pH 7.5), 150 mM NaCl, 1 mM PMSF, 5 mM MgCl2, and 25 mg DNaseI. 1% Lauryl Maltose Neopentyl Glycol (LMNG, Anatrace) was added to solubilize the membranes at 4 °C for 2 h under gentle agitation. After centrifugation (55,000g, 45 min, 4 °C) to remove the insoluble fraction, the supernatant containing the detergent-solubilized proteins was collected and loaded onto a column of CNBR- activated Sepharose beads (GE Healthcare) coupled to GFP nanobody (GFPnb). The GFPnb was derived from a llama single chain antibody and was expressed and purified as previously reported [37]. The columns were pre-equilibrated with buffer B [20 mM HEPES (pH 7.5), 150 mM NaCl, 0.1% LMNG, 0.01% cholesteryl hemisuccinate(CHS, Anatrace)]. After 1 h incubation at 4 °C, GFPnb resins were washed with 20 column volume of buffer B. The proteins were cleaved off the column with TEV protease at room temperature for 1 h. The TEV protease with a polyhistidine tag was removed by loading the supernatant to pre-equilibrated cobalt-based affinity resins (Talon, Clontech). The released proteins were collected, concentrated (Amicon 100 kDa cutoff, Millipore), and loaded onto a size-exclusion column (Superose 6 Increase 10/300 GL, GE Health Sciences) equilibrated with buffer C [20 mM HEPES (pH 7.5), 150 mM NaCl, 0.005% Glyco-diosgenin (GDN, Anatrace)]. The purified Heph protein was concentrated to ∼10 mg/mL, then immediately for making Cryo-EM sample, The rest of the protein was aliquoted and fast frozen in liquid nitrogen and stored at -80 °C for functional study.

## Cryo-EM sample preparation and data collection

Cryo grids were prepared on the Thermo Fisher Vitrobot Mark IV. Quantifoil R1.2/1.3 Cu grids were glow-discharged using the Pelco Easyglow. Concentrated Heph protein (3.5 μL) was applied to glow-discharged grids. After blotting with filter paper (Ted Pella) for 4.0-4.5 s, the grids were plunged into liquid ethane cooled with liquid nitrogen. For cryo-EM data collection, movie stacks were collected using EPU (Thermo Fisher Scientific) on a Titan Krios at 300 kV with a Quantum energy filter (Gatan), at a nominal magnification of ×105,000 and with defocus values of -2.2 to -0.8 μm. A K3 Summit direct electron detector (Gatan) was paired with the microscope. Each stack was collected in the super-resolution mode with an exposing time of 0.175 s per frame for a total of 50 frames. The dose was about 50 e- per Å2 for each stack. The movies were motion corrected with Relion 4 and binned (2 × 2)^24^ so that the pixel size was 0.832 Å and imported into CryoSPARC^25^ for further processing. Dose weighting was performed during motion correction, and the defocus values were estimated with Gctf^26^.

## Cryo-EM data processing

For full-length Heph dataset, 9674 micrographs were collected, of which 99 micrographs were excluded based on a CTF resolution limit of 6 Å, and a defocus limit of 30,000 Å. Initially, blob picking followed by two rounds of 2D classification were used to generate suitable templates for template picking, which then generated 7,776,718 particles. After three rounds of two- dimensional (2D) classification, 47 classes (containing 1,670,176 particles) were selected for ab initio three-dimensional (3D) reconstruction, which produced one good class with recognizable structural features with both soluble domain and transmembrane domain and two bad classes that did not have structural features. Both the good and bad classes were used as references in the heterogeneous refinement (cryoSPARC) and yielded a good class at 4.69 Å from 582,229 particles. Then nonuniform refinement (cryoSPARC) was performed with C2 symmetry and with masks for soluble domain and TM domain respectively, the soluble domain reaches overall resolution of 2.3 Å, the TM domain reaches a modest resolution around 6-8 Å which is able to recognize the helices.

Resolutions were estimated using the gold-standard Fourier shell correlationwith a 0.143 cut- off^27^ and high-resolution noise substitution^28^. Local res-olution was estimated using ResMap^29^.

## Model building and refinement

The structural models of Heph soluble domain were built based on the prediction model of AlphaFold2^30^, and sidechains were then adjusted based on the map. Model building was conducted in Coot^31^. Structural refinements were carried out in PHENIX in real space with secondary structure and geometry restraints^32^.

## Ferrozine assay and kinetic measurements of Heph activity

Heph ferroxidase activity was measured using a ferrozine (Fz) assay adapted from Deshpande et al. (2017)^15^, with modifications. Activity assays were performed at 37 °C in 100 mM Tris buffer (pH 6.8), and 0.005% GDN. Reaction mixtures (∼800 µL) contained 3.6 µM Heph and 50 µM Fe^2+^. At 15 s, 1.5 min, 5 min, and 8 min after reaction initiation, triplicate 45 µL aliquots were withdrawn and transferred to separate wells of a 96 well plate preloaded with 5 µL of 30 mM Fz. Addition of Fz chelated the remaining Fe^2+^ and quenched the reaction, yielding a final volume of 50 µL and a final Fz concentration of 3 mM per well. Absorbance of Fz-Fe^2+^ complex was measured at 570 nm using a BioTek spectrophotometer. Parallel reactions without Heph served as controls for nonenzymatic Fe²⁺ oxidation.

Before activity measurements, a standard curve was generated using defined Fe^2+^ concentrations prepared in the same assay buffer and mixed with Fz. The relationship between absorbance at 570 nm and Fe²⁺ concentration was fitted by linear regression and used to determine residual Fe²⁺ concentrations in the reaction samples. For D-penicillamine (DP) inhibition, Heph was incubated with 2 mM DP for 1 h during purification, immediately before the final size-exclusion chromatography step. The sample was subsequently loaded onto a Superose 6 Increase 10/300 GL column equilibrated with the standard SEC buffer containing 20 mM HEPES (pH 7.5), 150 mM NaCl, and 0.005% GDN to remove free DP. The activity of the resulting Heph preparation was measured using the Fz assay described above.

For kinetic measurements, reactions containing Fe^2+^ at initial concentrations of 0, 5, 10, 15, 25, or 75 µM were prepared in 100 mM Tris buffer (pH 6.8) at 37 °C. Reactions were initiated by adding Heph to a final concentration of 0.2 µM. At 15 s, 1, and 2 min, triplicate 45 µL aliquots were collected and analyzed using the same Fz procedure and standard curve described above. Initial reaction velocities were determined from the initial linear decrease in Fe^2+^ concentration over time. Initial velocities were plotted against the initial Fe^2+^ concentrations and fitted to the Michaelis-Menten equation by nonlinear regression in GraphPad Prism to determine Km and Vmax.

## NanoBiT luciferase complementation assay

Heph self-association in cells was assessed using a NanoBiT luciferase complementation assay^21,33^ (Promega, N2014). Full-length human Heph was fused at its C-terminus to either the large (LgBiT) or small (SmBiT) NanoBiT fragment, generating Heph-LgBiT and Heph-SmBiT, respectively.

HEK293S GnTI⁻ cells were seeded in 96-well plates and cotransfected with plasmids encoding Heph-LgBiT 2 ng per well and Heph-SmBiT 2 ng per well using Lipofectamine 3000 (Thermo Fisher Scientific). At 48 h after transfection, luminescence was measured using Nano-Glo Live Cell Reagent, prepared from the Nano-Glo Live Cell Substrate and LCS Dilution Buffer supplied with the kit. Luminescence was measured using a BioTek microplate reader. Cells coexpressing the kit-supplied SmBiT-PRKACA and LgBiT-PRKAR2A constructs served as the positive control. Cells coexpressing Heph-LgBiT and the kit-supplied HaloTag-SmBiT construct served as the negative control. Data are presented as mean ± SD (n = 3 per group). Statistical analyses were performed in GraphPad Prism using ordinary one-way ANOVA followed by Tukey’s multiple comparisons test. Adjusted P values < 0.05 were considered statistically significant.

## Data Availability

The atomic coordinates of human Heph have been deposited in the Protein Data Bank (http://www.rcsb.org) under the accession code 39EP. Its corresponding electron microscopy map has been deposited in the Electron Microscopy Data Bank (https://www.ebi.ac.uk/pdbe/emdb/) under the accession code EMD-79355. Source data are provided with this paper.

## Acknowledgements

This work was supported by grants from National Institute of Health (HL157473 to Y.P., DK122784, GM145416, and GM151548 to M.Z.). We acknowledge the cryo-EM core at Baylor College of Medicine, which is supported in part by CPRIT Core Facility Award RP190602. We thank Dr. Gaya P. Yadav for cryo-EM data collection at the Laboratory for Biomolecular Structure and Dynamics (LBSD) of Texas A&M University. The LBSD is supported, in part, by the Department of Biochemistry and Biophysics, AgriLife, Texas A&M University, and CPRIT Facility Support Grant (RP25049).

## Notes

### Competing Interest Statement

The authors have declared no competing interest.

