## Supplemental Figure1-3; Table1 for "Structure of human hephaestin reveals a homodimer anchored to the cell membrane"

**Supplementary Figures:**

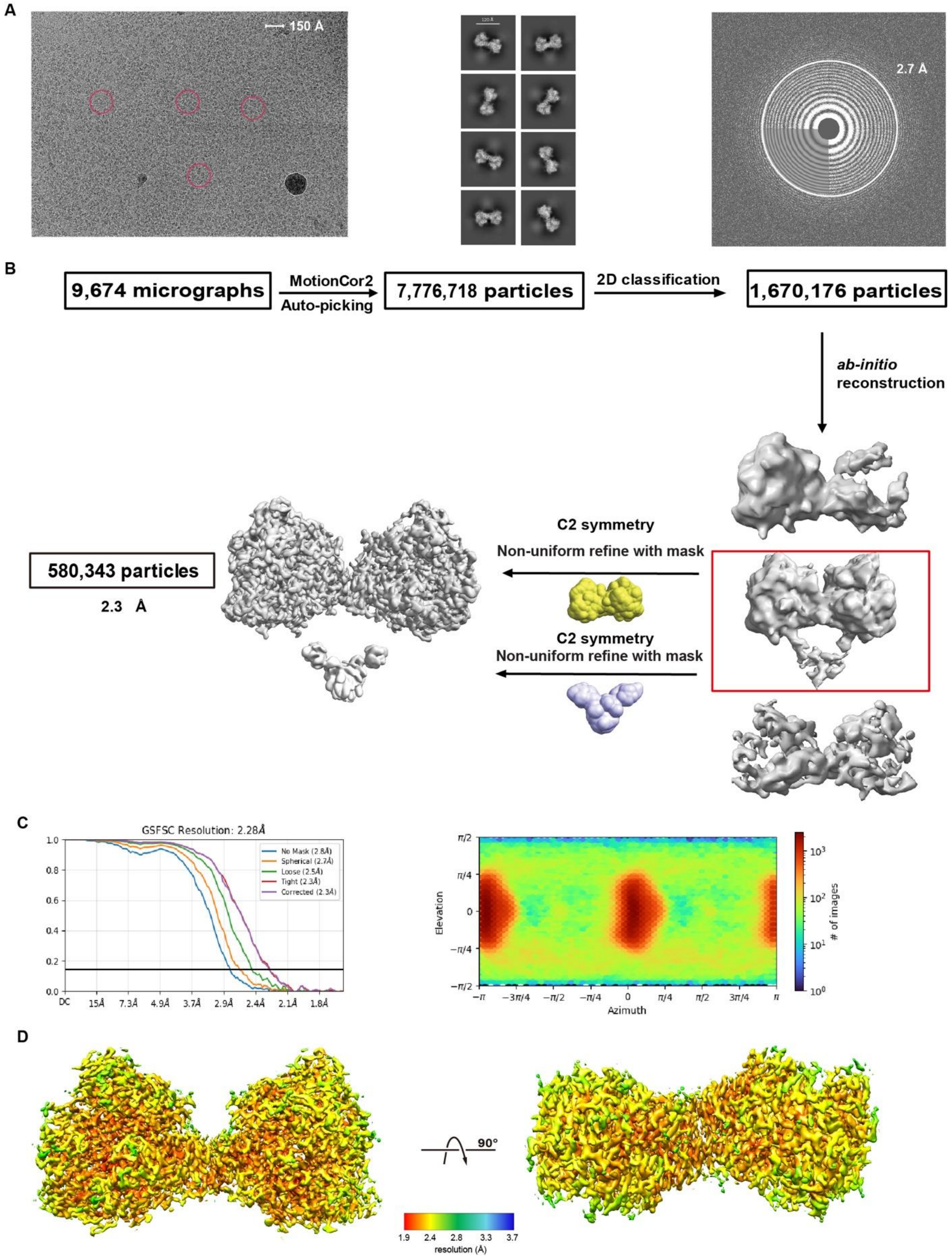


**Supplementary Figure 1. Cryo-EM data processing and validation of the human Heph reconstruction.** Representative cryo-EM micrograph, with selected Heph particles marked by red circles (left), representative two-dimensional class averages (middle), and a power spectrum showing Thon rings (right). Scale bar, 150 Å. (B) Overview of particle selection, classification, and three-dimensional reconstruction. The selected ab initio class is outlined in red. Separate masks encompassing the extracellular region (yellow) and transmembrane region (purple) were used for nonuniform refinement with C2 symmetry. The extracellular reconstruction reached an overall resolution of approximately 2.3 Å. Particle numbers at the indicated processing stages are shown. (C) Gold-standard Fourier shell correlation (FSC) curves (left) and particle angular distribution (right) for the final extracellular reconstruction. Resolution was determined using an FSC threshold of 0.143. (D) Two views of the extracellular density map, related by a 90° rotation and colored by local resolution according to the accompanying scale.

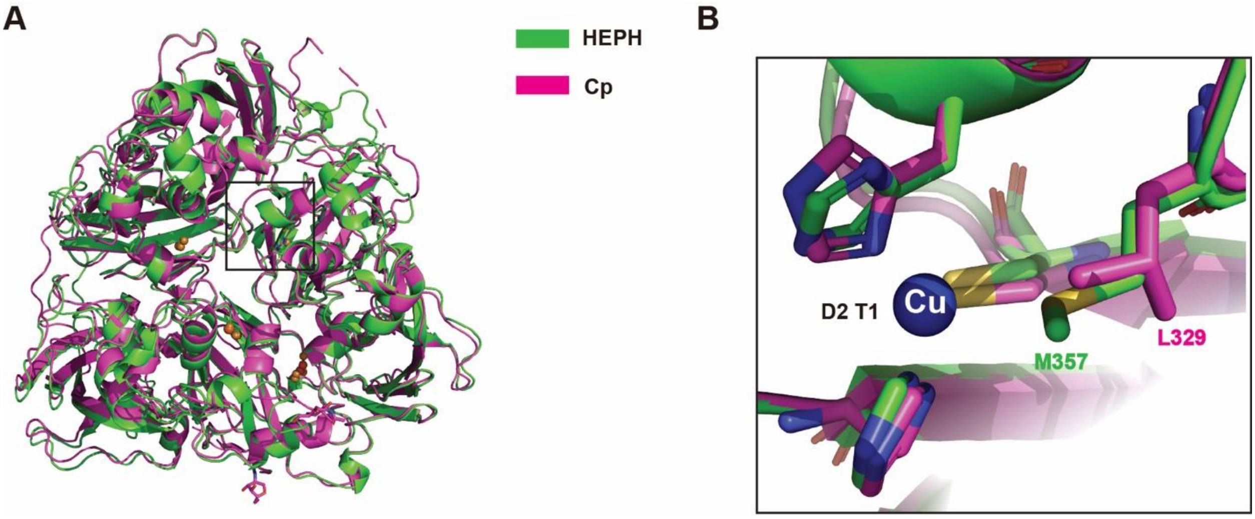


**Supplementary Figure 2. Structural comparison of Heph and ceruloplasmin.** (A) Superposition of Heph (green) and ceruloplasmin (magenta; PDB: 1KCW), yielding a root-mean-square deviation (RMSD) of 2.0 Å across 1,002 aligned pairs. The boxed region is enlarged in panel B. (B) Close-up of the domain 2 type 1 (D2 T1) copper-binding site, highlighting Met357 in Heph and the corresponding Leu329 in ceruloplasmin. Side chains are shown as sticks. The copper ion (Cu) is shown as a blue sphere.

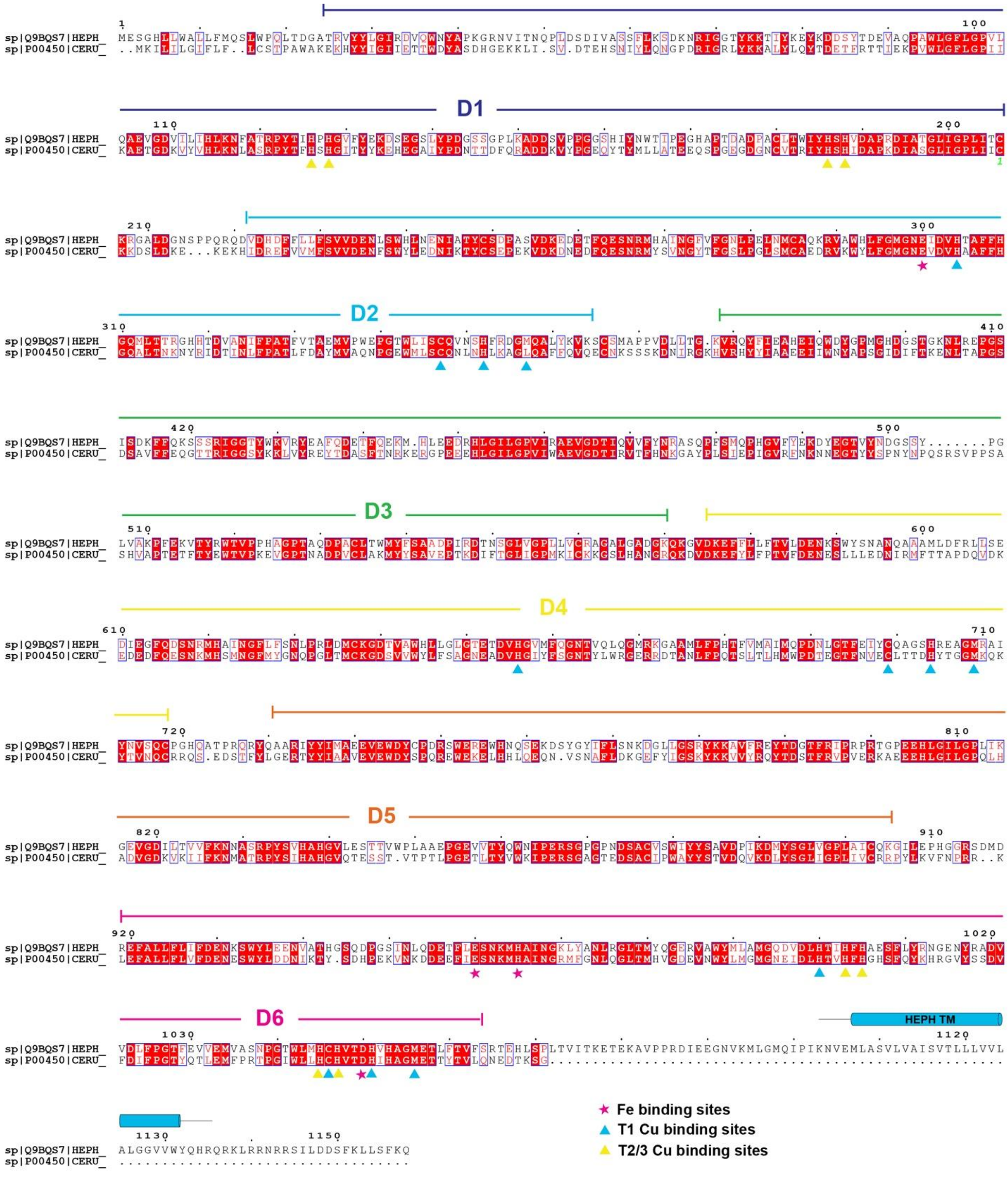

**Supplementary Figure 3. Sequence alignment of human Heph and ceruloplasmin.** The amino acid sequences of human Heph (UniProt Q9BQS7) and ceruloplasmin (UniProt P00450) were aligned using the Clustal Omega server[34] and visualized using the ESPript server[35]. Colored lines above the alignment delineate the six cupredoxin-like domains of Heph (D1–D6). The Heph transmembrane segment is indicated by a cyan bar labeled ‘HEPH TM’. Magenta stars mark iron-binding residues, cyan triangles mark type 1 copper-binding residues, and yellow triangles mark type 2 and type 3 copper-binding residues.

**Supplementary Table 1. Summary of cryo-EM data collection, processing, and structural refinement**

| | Human hephaestin (EMD-79355, PDB 39EP) |
|---|---|
| **Data collection and processing** | |
| Magnification | 105,000 |
| Voltage (kV) | 300 |
| Electron exposure (e–/Å$^2$) | 50 |
| Defocus range (μm) | [-2.0, -0.8] |
| Pixel size (Å) | 0.832 |
| Symmetry imposed | C2 |
| Initial particle images (no.) | 7,776,718 |
| Final particle images (no.) | 580,343 |
| Map resolution (Å) | 2.3 |
| FSC threshold | 0.143 |
| Map resolution range (Å) | 1.9-3.7 |
| **Refinement** | |
| Initial model used (PDB code) | Alphafold2 |
| Model resolution (Å) | 2.4 |
| FSC threshold | 0.5 |
| Map sharpening *B* factor (Å$^2$) | -87 |
| Model composition | |
| Non-hydrogen atoms | 16,596 |
| Protein residues | 2086 |
| Ligands | NAG:2, OXY:2, FE2:2, CU:12 |
| *B* factors (Å$^2$) | |
| Protein | 35.7 |
| Ligand | 52.8 |
| R.m.s. deviations | |
| Bond lengths (Å) | 0.003 |
| Bond angles (°) | 0.539 |
| Validation | |
| MolProbity score | 0.8 |
| Clashscore | 1.02 |
| Poor rotamers (%) | 0.79 |
| Ramachandran plot | |
| Favored (%) | 98.56 |
| Allowed (%) | 1.44 |
| Disallowed (%) | 0 |
